# Viral proteins and virus-like particles of the LTR5_Hs endogenous retrovirus in human primordial germ cell-like cells

**DOI:** 10.1101/2022.09.24.509338

**Authors:** Mutsumi Kobayashi, Misato Kobayashi, Johannes Kreuzer, Eric Zaniewski, Jae Jung Kim, Keiko Shioda, Hikari Hagihara, Junko Odajima, Ayako Nakashoji, Yi Zheng, Jianping Fu, Maria Ericsson, Kazuhiro Kawamura, Shannon L. Stott, Daniel Irimia, Wilhelm Haas, Chin-Lee Wu, Maria Tokuyama, Toshi Shioda

## Abstract

The hominoid-specific endogenous retrovirus LTR5_Hs is transcriptionally activated in human primordial germ cell-like cells (hPGCLCs), a pluripotent stem cell-derived cell culture model of PGCs. Here, taking the unique advantage of our novel cell culture method to obtain large amounts of pure hPGCLCs, we performed proteomics profiling of hPGCLCs and detected various viral proteins produced from the LTR5_Hs RNA *via* ribosomal frameshifting. We also present transmission electron microscopy images of 100-nm diameter virus-like particles (VLPs) assembled at the surface of hPGCLCs. Compared to hPGCLCs, expression of LTR5_Hs RNA is far weaker in human seminomas, the germ cell tumors resembling PGCs. Re-analysis of published single cell RNA-seq data of human embryos revealed strong activation of LTR5_Hs in migrating PGCs but suppressed in PGCs upon they reach the gonadal anlagen. In the microfluidics-supported polarized embryoids mimicking peri-implantation stages of human embryos, LTR5_Hs RNA was detected by RNA in situ hybridization in NANOG^+^/TFAP2C^+^/SOX17^+^ cells resembling freshly emerged PGCs. These results support that human germ cells produce LTR5_Hs proteins and VLPs during their earliest stages of normal development until their settlement in the gonadal anlagen.

**SUMMARY STATEMENT:** The hominoid-specific endogenous retrovirus LTR5_Hs is activated in a cell culture model resembling early-stage human primordial germ cells, producing not only viral RNA but also retrovirus proteins and virus-like particles.

## INTRODUCTION

Human endogenous retroviruses (HERVs) are remnants of ancient infection by retroviruses, comprising nearly 8% of the human genome (Deniz et al., 2018; Durnaoglu et al., 2021a; Geis and Goff, 2020; Groh and Schotta, 2017; Liu et al., 2014; Mao et al., 2021). Whereas most HERVs are permanently inactivated by accumulated mutations or strongly suppressed by epigenetic machineries, some of the activation-competent copies of HERVs may play critical roles in a wide variety of human diseases, including various malignancies, autoimmune diseases, and neurological disorders (Babaian and Mager, 2016; Doucet-O’Hare et al., 2021). Non-physiological reactivation of HERVs may be caused by malnutrition, exposure to environmental toxicants, or impaired health conditions (Sakurai et al., 2019; Sharif et al., 2013; Shioda et al., 2022). On the other hand, activation of several copies of HERVs is necessary for normal human development or body functions (Evsikov and Marin de Evsikova, 2016; Weiss, 2016). For example, production of the salivary alpha-amylase (ptyalin) is dependent on salivary gland-specific activation of a copy of HERV located in the promoter of the *AMY1C* gene (Ting et al., 1992). Formation of the syncytiotrophoblast through trophoblast cell fusion in placenta requires syntytin-1, a fusogenic protein derived from the envelope protein of a HERV (Durnaoglu et al., 2021b; Mi et al., 2000). Strong transcriptional activation of the HERV-H family is necessary for acquisition and maintenance of pluripotency by stem cells during early development of human embryos, possibly through affecting the chromatin contact dynamics (Ohnuki et al., 2014; Sexton et al., 2022; Zhang et al., 2019).

Members of the HERV-K clade represent the HERVs most recently integrated into the human genome, and many of them are still transcriptionally active (Garcia-Montojo et al., 2018; Xue et al., 2020a). HERV-K consists of ten or eleven HML (human mouse mammary tumor virus-like) subgroups, among which HML-2 is the youngest and most active (Subramanian et al., 2011). Although all copies of the HML-2 proviruses are defective in at least one gene, many of them have complete open reading frames encoding retroviral proteins detected in healthy and malignant human cells (Curty et al., 2020). HML-2 are capable of forming virus-like particles (VLPs), which has been detected in human naïve pluripotent stem cells as well as malignant cells (Bieda et al., 2001; Grow et al., 2015). Whereas the human genome contains approximately 1000 copies of HML-2 solitary LTRs, which lack DNA sequences coding viral proteins, only less than 100 copies of HML-2 proviruses have been identified so far (Xue et al., 2020b). The HML-2 family of HERVs consists of three subgroups – namely, LTR5_Hs, LTR5A, and LTR5B. LTR5_Hs is the youngest among all types of HERVs and has successfully expanded in the hominoid lineage (Garcia-Montojo et al., 2018; Holloway et al., 2019; Xue et al., 2020a). LTR5_Hs is activated in early-stage pluripotent cells in human embryos and embryonal carcinoma tumor cells, in which their transcriptional actions significantly affect the epigenomic integrity of the human genome (Fuentes et al., 2018; Grow et al., 2015; Pontis et al., 2019; Zhang et al., 2022).

Human Primordial Germ Cells (hPGCs) emerge from amnion and epiblast of embryos as the earliest precursors of all germ cells 11-12 days after fertilization (Saitou, 2021). Despite the importance of studying hPGCs to promote reproductive health, access to hPGCs in human embryos is extremely challenging for both technical and ethical barriers. To overcome these hurdles, cell culture models resembling hPGCs have been generated from human pluripotent stem cells (hPSCs) (Saitou, 2021). These models, collectively known as human PGC-Like Cells (hPGCLCs), can be produced from various states of naïve pluripotent stem cells (Irie et al., 2015; Mitsunaga et al., 2017; von Meyenn et al., 2016) or cells resembling the early-stage mesodermal precursor cells (incipient Mesoderm-Like Cells: iMeLCs) (Chen et al., 2017; Sasaki et al., 2015). Our previous study showed that the transcriptomic profiles of hPGCLCs produced using various methods in different laboratories are largely homogenous, resembling the transcriptome of hPGCs before initiation of the chemotaxis towards gonadal anlagen (Mitsunaga et al., 2017). hPGCLCs are capable of differentiating to advanced stages of male and female germ cells *in vitro*, further demonstrating their faithful resemblance to hPGCs (Hwang et al., 2020; Yamashiro et al., 2018). Whereas *in vitro* expansion of hPGCLCs has been proven challenging (Gell et al., 2020; Murase et al., 2020), our recent study has overcome this technical barrier and established a serum-free, feeder layer-free cell culture condition that effectively supports long-term expansion of hPGCLCs (Kobayashi et al., 2022). Under this condition, Long-Term Culture hPGCLCs (LTC-hPGCLCs) strongly express telomerase and rapidly amplify without apparent passaging limit or signs of senescence while strictly maintaining their hPGC-like characteristics as a highly homogeneous cell population. LTC-hPGCLCs provide unprecedented opportunities to obtain large amounts of pure hPGCLC specimens, which are often required for several standard analytical approaches such as proteomics or transmission electron microscopy (TEM) (Graham and Orenstein, 2007).

Recent studies have shown that LTR5_Hs are activated in hPGCLCs and provided evidence that this hominoid-specific group of the HERVs play significant roles in transcriptional regulation of genes involved in development of germ cells (Ito et al., 2022; Xiang et al., 2022). Our study has shown specific and robust CpG demethylation of LTR5_Hs in both fresh and long-term cultured hPGCLCs compared to the precursor hiPSCs (Kobayashi et al., 2022). In freshly isolated hPGCLCs, less than 20% of CpG sites in LTR5_Hs were methylated whereas other HERVs such as LTR7/HERV-H retained nearly 50% CpG methylation. Long-term expansion of hPGCLCs for 12 weeks further reduced CpG methylation in LTR5_Hs down to ∼10%. Thus, activation of LTR5_Hs in hPGCLCs is specific – it is not a mere consequence of global DNA demethylation in this model of human germ cells.

Taking advantage of the LTC-hPGCLCs, our current study demonstrates that not only LTR5_Hs viral RNA species but also various retroviral proteins produced by the ribosomal frameshifting are strongly expressed in this cell culture model resembling early-stage normal hPGCs. In contrast, expression of LTR5_Hs RNA in human seminomas, which are derived from transformed PGCs and still expressing the PGC marker SOX17 (Muller et al., 2021), is proven weak. We also show TEM images capturing robust assembly of LTR5_Hs VLPs at the plasma membrane of LTC-hPGCLCs. Using an *in vitro* model resembling the peri-implantation stages of human embryos formed under a condition of microfluidics-aided polarized exposure to bone morphogenetic protein 4 (BMP4), we present evidence that activation of LTR5_Hs occurs as soon as hPGCs emerge from their precursors. Thus, our study provide evidence that the earliest stages of normal human germ cell development – from the germline specification to hPGC settlement in the gonadal anlagen – occurs in the presence of various retrovirus-like activities of LTR5_Hs, involving not only their transcriptional actions but also production of various retroviral proteins. Our study also suggests that hPGCs may robustly produce VLPs and deposit the particles in the path of their migration.

## RESULTS

### RNA expression from distinct groups of HERVs in human iPSCs (hiPSCs), hPGCLCs, non-germline human embryoid body cells (hEBCs), and human seminoma tumors

Recent studies revealed strong activation of the youngest HERV species LTR5_Hs in hPGCLCs (Ito et al., 2022; Xiang et al., 2022) whereas an older HERV LTR7/HERV-H are robustly activated in their precursor hPSCs (Ohnuki et al., 2014; Sexton et al., 2022; Zhang et al., 2019). Using the ERVmap tool of quantitative determination of RNA expression from HERV loci (Tokuyama et al., 2018) and RNA-seq data we previously published (Mitsunaga et al., 2017), we examined HERV RNA expression in the CD38-positive hPGCLCs, their precursor hiPSCs (clones A4, A5, A6), and CD38-negative non-germline cells. To this analysis we also included total RNA specimens isolated from ten cases of human pure seminomas, which are transformed late-stage hPGCs (Oosterhuis and Looijenga, 2019).

Unsupervised hierarchical clustering successfully classified the specimens by cell/tissue types – namely, hiPSCs, hPGCLCs, hEBCs, and seminomas – based solely on expression of HERV RNA transcripts (Fig. 1A, *left* heatmap), reproducing our previous analysis using the whole transcriptomes of protein coding genes (Mitsunaga et al., 2017). Ten clusters of HERVs differentially expressed between distinct types (C1-C10) were identified (Fig. 1A, connecting *left* and *right* heatmaps, and Table S1). Clusters C3 and C6 consisted of two subclusters (C3a and C3b, C6a and C6b) located separately in the main (*left*) heatmap. Relative expression profile of HERVs representing each of these ten clusters across different cell/tissue types demonstrated striking type-specific expression of HERVs (Fig. 1B). Agreeing with previous studies, hiPSCs strongly expressed LTR7/HERV-H, and the majority of HERVs specifically expressed in hiPSCs (Cluster 2) were LTR7/HERV-H, which was also the dominant HERV species commonly expressed in both hiPSCs and seminomas (Cluster 1) or hiPSCs and hPGCLCs (Cluster 3) (Fig. S1, Table S1). In contrast, among 96 HERVs specifically expressed in hPGCLCs (Cluster 4), LTR5_Hs was the most frequently found HERV species over LTR7/HERV-H. Among 32 HERVs specifically expressed in seminomas (Cluster 10), we detected only one or three copies of LTR7/HERVH-int or LTR5_Hs, respectively, whereas LTR17/LTR17-int was the most frequently activated HERV species. We identified only 9 HERVs commonly activated in both hPGCLCs and seminomas (Cluster 9), and none of them was LTR5 and only one was LTR5/HERV-H. These results showed that LTR7/HERV-H represented HERVs activated in hiPSCs. Upon differentiation of hiPSCs to hPGCLCs, LTR5_Hs was activated while LTR7/HERV-H was suppressed. LTR5_Hs activation was not significant in seminoma tissues.

**Fig. 1.**
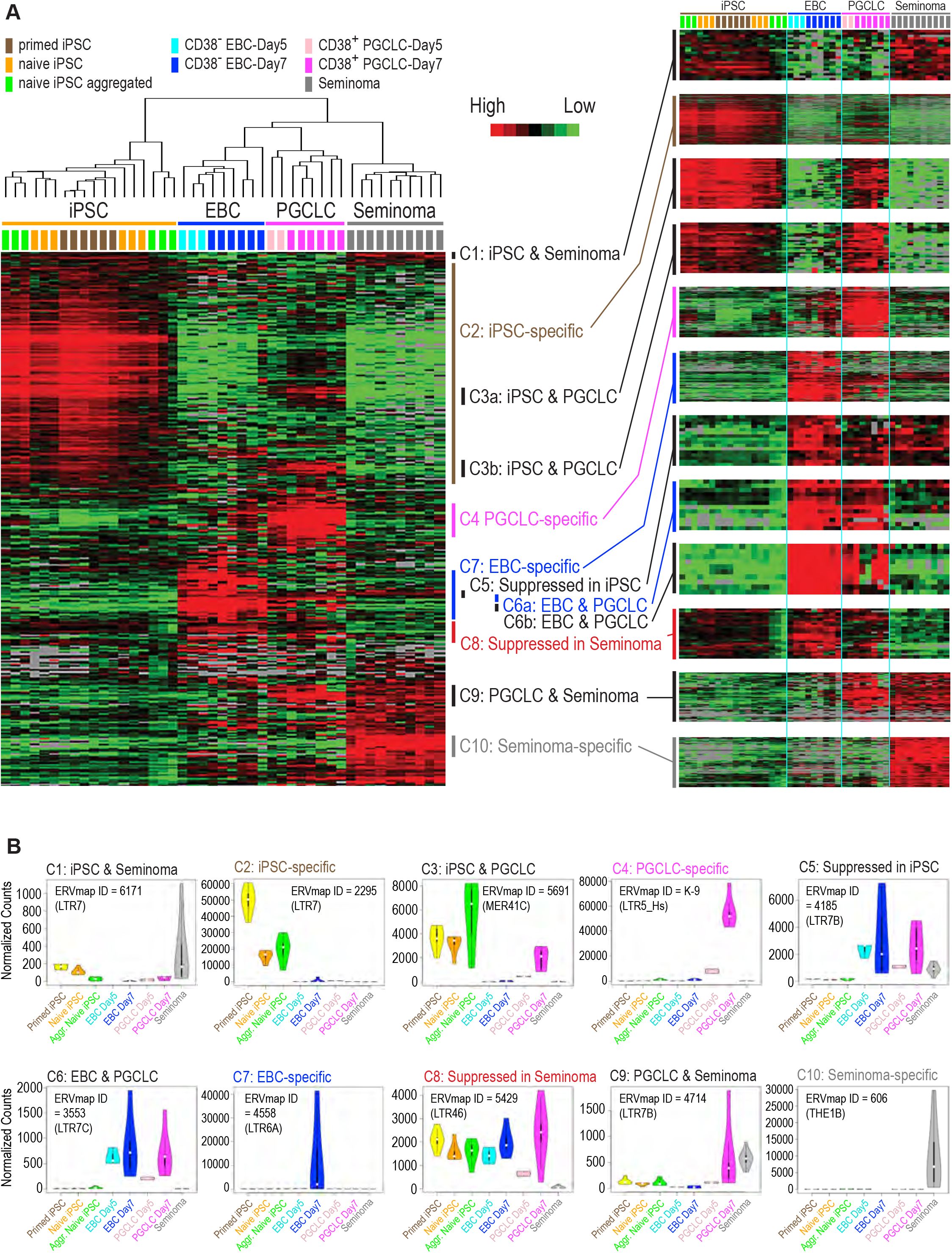
RNA-seq profiling of human iPSCs, embryoid bodies, PGCLCs, and seminoma tissues for expression of HERV RNA using ERVmap. (A) Heatmap representations of unsupervised clustering of HERV RNA expression. Color-coded cell/tissue types are shown on top of each heatmap. Clusters of HERVs identified for the left heatmap are magnified in the right heatmap. (B) RNA expression of HERVs representing each cluster across eight cell/tissue types. Normalized RNA counts are shown in violin plots. Point, bar, and whiskers of the boxplot part in the violin shape indicate median, Q1 and Q3 quartiles, and minimum/maximum values. ERVmap IDs of HERVs and their clades (in parentheses) are shown for each panel.

We determined 50 copies of HERVs most strongly expressed in hPGCLCs, and we summarized their locations in the human genomic DNA and strength of viral RNA expression in Table S2. Among them, 40 copies (80%) belonged to Cluster 4 (specific to PGCLCs) whereas 7 copies (14%) belonged to Cluster 9 (PGCLCs and seminomas). Among these 40 Cluster 4 HERVs, 20 copies (50%) were LTR5, and all of them were LTR5_Hs. In contrast, 2 copies of the Cluster 9 HERVs were LTR5, and one of them was LTR5_Hs. Thus, HERVs strongly activated specifically in hPGCLCs were represented by LTR5_Hs.

### Evaluation of computational tools for quantitative determination of HERV viral RNA expression from RNA-seq data

ERVmap is a software tool developed for quantitative analysis of RNA-seq data for expression of viral RNA transcripts from HERVs (Tokuyama et al., 2018). Several other computational tools for similar purposes have been described, but accuracy of these tool is a debatable subject (Iniguez et al., 2019; Tokuyama et al., 2019). To establish a reliable computational pipeline for HERV RNA expression, we compared representative tools – namely, ERVmap (Tokuyama et al., 2018), Telescope (Bendall et al., 2019), and Salmon-TE (Jeong et al., 2018).

The original ERVmap consist of a series of Perl script and requires several components that are no longer available from open sources. We re-implemented ERVmap using the scripting language Ruby and open-source codes to create ERVmap2. Whereas ERVmap assigns RNA-seq reads to 3,220 hand-picked HERV proviruses in the GRCh38/hg38 human reference genome, we generated an independent list of relatively well-integrated 2,504 HERV proviruses consisting of one 5’ LTR, one 3’ LTR, and at least one internal sequence connected via gaps not greater than 1 kb (Fig. S1A). Numbers of HERVs belonging to each clade and the whole list of the selected HERVs (which is referred to as the ERVmap2 HERV provirus list in this study) are provided as Tables S3 and S4, respectively. The majority of the selected, well-organized HERV proviruses are HERV-H (37%), HERV-L (20%), or HERV9 (10%); only 55 copies (2.2%) of HML2 proviruses, including LTR5_Hs, were included in this list (Fig. S1B).

From the ERVmap2 list (BED format) or its GTF-format version required for Telescope, we generated DNA sequences of well-organized HERV proviruses in the FASTA format (Fig. S2A). Using the ART simulator of Illumina sequencing data (Huang et al., 2012) and these FASTA provirus sequences, we generated “gold standard” SAM alignment data and FASTQ simulated reads. The simulated FASTQ reads were then supplied to ERVmap, ERVmap2, Telescope, or Salmon-TE to estimate normalized expression of HERV RNA transcripts. On the other hand, HERV RNA expression levels were calculated directly from the gold standard SAM data and compared with the outcomes of the above tools by X-Y hexagon plots, in which a greater correlation coefficient reflects a greater degree of accuracy (Figs. S2B-S2E). Among these tools, ERVmap2 showed the greatest level of accuracy (*R*^*2*^ = 0.8687; Fig. S2C) followed by Salmon-TE (*R*^*2*^ = 0.7655; Fig. S2E) and ERVmap (*R*^*2*^ = 0.5707; Fig. S2B). Note that the same number of datum points were plotted in each panel although highly overlapped points reduce numbers of visible points. Whereas ERVmap2 and Salmon-TE over- and under-estimate HERV RNA expression relatively evenly, ERVmap tended to be biased toward under-estimation. On the other hand, the correlation coefficient of Telescope (*R*^*2*^ = 0.002488; Fig. S2D) was significantly lower than those of other tools with strong over- and under-estimation of HERV RNA expression. When the hierarchical clustering analysis shown in Fig. 1A was performed using Telescope, RNA specimens were classified by their types with significantly reduced accuracy, and identification of type-specific HERV clusters was practically challenging (Fig. S3). These results support that ERVmap2 is an adequate tool for quantitative evaluation of RNA transcripts from HERV proviruses.

### HERVH-to-HML2 class switching in HERV RNA expression during hiPSC differentiation to hPGCLC

Taking advantage of the accurate detection of HERV RNA from RNA-seq data implemented by ERVmap2, we determined relative amounts of viral RNA transcripts expressed from the 18 clades of HERVs defined in Table S3. RNA of LTR7/HERV-H was very strongly expressed in hiPSCs but significantly suppressed in hPGCLCs (Fig. 2A). In contrast, HML2 RNA was strongly expressed in hPGCLCs whereas it was nearly undetectable in primed hiPSCs. Expression of viral RNA from other 16 clades was far weaker than the above two clades in hiPSCs or hPGCLCs.

**Fig. 2.**
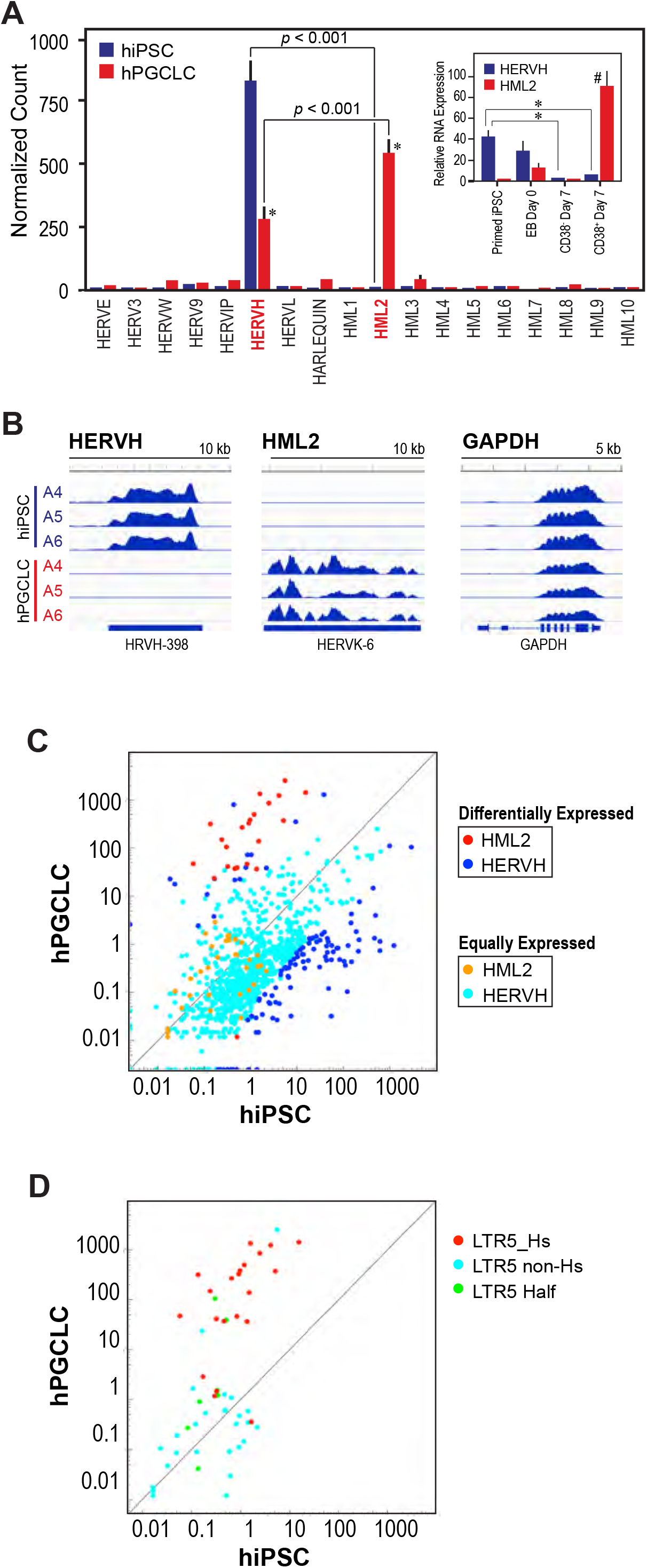
Activation of the LTR5_Hs human-specific HERVs in PGCLCs. (A) Expression profiles of HERV RNA in human iPSCs and PGCLCs. Normalized RNA expression of HERVs was calculated from RNA-seq data of primed human iPSCs and hPGCLCs using ERVmap2 and presented for 18 HERV clades. Asterisk indicates statistical significance (*p* < 0.01) between iPSC and PGCLC in each clade. *Inset:* Reverse transcription qPCR determination of RNA expression from HERVH and HML2 in primed iPSCs, embryoid body (EB) cells at day 0 culture, CD38^+^ PGCLCs at day 7 culture, and CD38^-^ EB cells at day 7 culture. Relative amounts of RNA to those in CD38^-^ EB cells (defined as 1) are shown. Sharp indicates statistical significance (*p* < 0.01) to each of other cell types. Asterisk indicates significance (*p* < 0.01). For both the main and inlet panels, each bar shows mean ± SEM of data obtained from three independent human iPSC clones and PGCLCs generated from them. (B) Representative RNA-seq tracks of three independent human iPSC clones (A4, A5, A6) and PGCLCs derived from them for HERVH, HML2, and GAPDH. The bigWig tracks are normalized for each RNA for direct comparisons across all 6 RNA-seq data. (C) Differential expression of individual copies of HERVH and HML2 between human iPSCs and PGCLCs. Scatter plots shows statistically significant and insignificant differential expression as indicated. (D) Differential expression of HML2 species LTR5_Hs, non-Hs LTR5, and LTR5 Half copies between human iPSCs and PGCLCs.

Real-time qPCR quantitation has successfully verified the ERVmap2 quantitation of LTR7/HERV-H and HML2 viral RNA transcripts (Fig. 2A *inset*). Expression of LTR7/HERV-H RNA was already diminished in the naïve hiPSCs comprising the freshly formed embryoid bodies (EB Day 0) compared to the primed hiPSCs. On the other hand, expression of HML2 was very weak in the primed hiPSCs but already augmented in the naïve hiPSCs (EB Day 0). After 7 days of incubation of the embryoid bodies, HML2 RNA was strongly expressed in the CD38^+^ hPGCLCs but suppressed in the CD38^-^ non-germline cells to a nearly undetectable level. Thus, the classes of strongly activated HERVs are switched from LTR7/HERV-H to HML2 during the conversion of primed hiPSCs to hPGCLCs. Typical RNA-seq tracks demonstrating this class switching are shown in Fig. 2B.

We next examined the relative strength of RNA expression between HERVs differentially or equally expressed in hiPSCs and hPGCLCs (Fig. 2C). Amounts of RNA expressed from differentially expressed copies of LTR7/HERV-H (*pale blue dots*) are largely comparable to those of equally expressed copies (*dark blue dots*). In contrast, HML2 expression from differentially expressed copies (*red dots*) were stronger than those of equally expressed copies (*yellow dots*). The apparent absence of HML2 copies strongly expressed in both hiPSC and hPGCLCs suggests that HML2 is actively suppressed in hiPSCs.

In our ERVmap2 analysis of well-organized HERVs, each copy of HML2 has 5’- and 3’- end LTRs belonging to three subclasses of LTR5 – namely, LTR5_Hs, LTR5_A, and LTR5_B. Some of the HML2 copies have two LTR5_Hs at both end whereas other copies may contain one or two non-Hs LTR5. The majority of the HML2 copies strongly expressed in hPGCLCs exclusively possessed LTR5_Hs (Fig. 2D, LTR5_Hs) whereas most HML copies harboring one or two non-Hs LTR5 (LTR5 Half or LTR5 non-Hs, respectively) were expressed in both hiPSC and hPGCLCs but very weakly.

### Expression of HML2 HERV RNA in early-stage PGCs *in vivo*

Since hPGCLCs resembles early-stage, DAZL-negative hPGCs in human embryos at 8-weeks of gestation or earlier (Hwang et al., 2020; Kobayashi et al., 2022; Mitsunaga et al., 2017), we attempted to detect HML2 viral RNA in previously published single-cell RNAseq data of human male and female germ cells at 4-26 weeks of gestation (Li et al., 2017). tSNE plots clearly separated NANOG^+^ sexually bipotential germ cells from sexually committed germ cells, including SIX1^+^ male cells and cells expressing female germline markers STRA8, SYCP1, and ZP3 (Figs. 3A and 3B). Cells strongly expressing HML2 were DAZL^-^, 4-5 weeks germ cells in both male and female embryos. Modest expression of HML2 was also observed with NANOG^+^/DAZL^+^ immature germ cells. On the other hand, HERV-H RNA was weaklly expressed in all stages of germ cells. Expression of HML2 was stronger in mitotic germ cells of both male and female embryos whereas HERV-H was expressed equally in all stages of germ cells except for strong expression in female post-meiotic cells (Fig. 3C). These data indicate that HML2 is strongly activated in early-stage PGCs in 4-5 weeks embryos and thereafter suppressed upon sexual differentiation of germ cells.

**Fig. 3.**
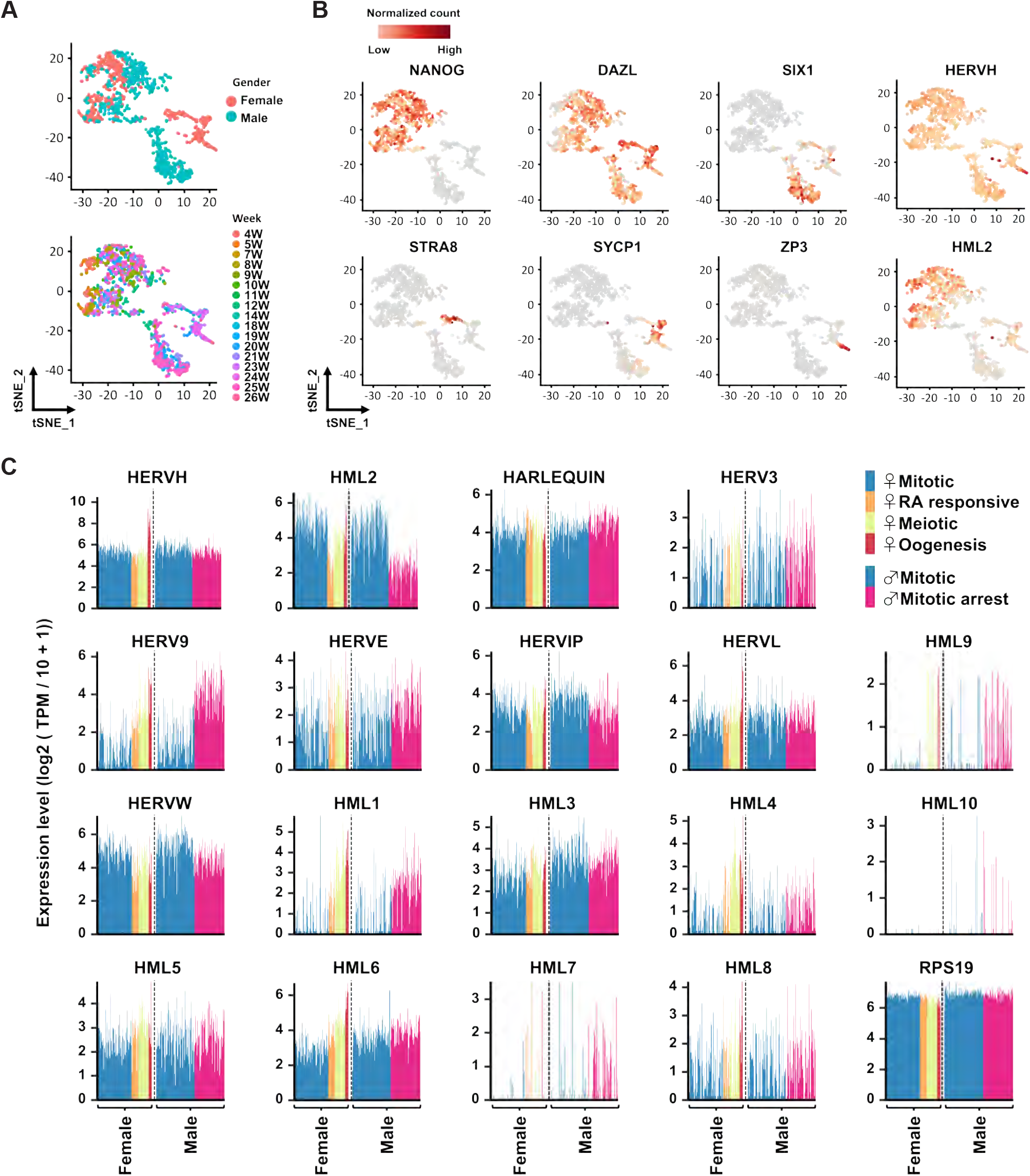
Expression of HERV RNA in human fetal germ cells. (A, B) Single cell RNA-seq data of human fetal germ cells (Li, 2017) are presented as tSNE plots with color indexes for sex (A, *top*), gestational weeks (A, *bottom*), and marker genes (B). (C) Expression of 18-clades of HERV RNA in fetal germ cells at various developmental stages. Each subpanel shows relative expression of a HERV RNA species in female and male gonads at developmental stages color-coded as indicated. TPM, transcripts per kilobase million

### Activation of HML2 immediately after hiPSC differentiation to hPGCLCs

hPGCs emerge from amnion and epiblast of embryos 11-12 days after fertilization (Saitou, 2021). In human embryos at 4-weeks of gestation, migrating hPGCs already express large amounts of HML2 viral RNA (Fig. 3). To estimate the timing of HML2 activation during the very early stages of human germ cell development, we took advantage of the microfluidics-supported, hPSC-derived human embryoid model that recapitulates critical landmarks of pre-gastrulation development under polarized exposure to BMP4 (Zheng et al., 2019). In this model, aggregates of hPSCs are formed in slits connecting two microfluidics channels, one of which is filled with gel (Gel channel) for physical support of the aggregates and the other (Cell-loading channel) is used for polarized supply of BMP4 as well as loading cells to the slits (Fig. 4A). In the absence or presence of polarized BMP4, the aggregates grew to epiblast-like cysts (ELCs) or posteriorized embryonic-like sac (P-ELS), respectively (Fig. 4B). In both ELCs and P-ELSs, NANOG was expressed in the epithelial parts of the embryoids (Fig. 4C). PGC-like cells emerged as NANOG/TFAP2C/SOX17 triple-positive cells in P-ELSs but not in ELCs (Fig. 4C), reproducing the original study of this model (Zheng et al., 2019). RNA *in situ* hybridization (RNA-ish) of P-ELSs detected HML2 viral RNA exclusively in the PGC-like cells expressing nuclear SOX17 protein, and all these SOX17-expressing cells are HML2 viral RNA-positive (Fig. 4D). In contrast, no HML2 RNA-ish signal was detected in ELCs (data not shown). Simultaneous RNA-ish detection of NANOG and HML2 RNA revealed that HML2-positive NANOG-positive are mostly overlapped (Fig. 4E). These results provide *in vitro* evidence that HML2 is specifically activated in hPGCs immediately after they emerge in amnion/epiblast of pre-gastrulation human embryos.

**Fig. 4.**
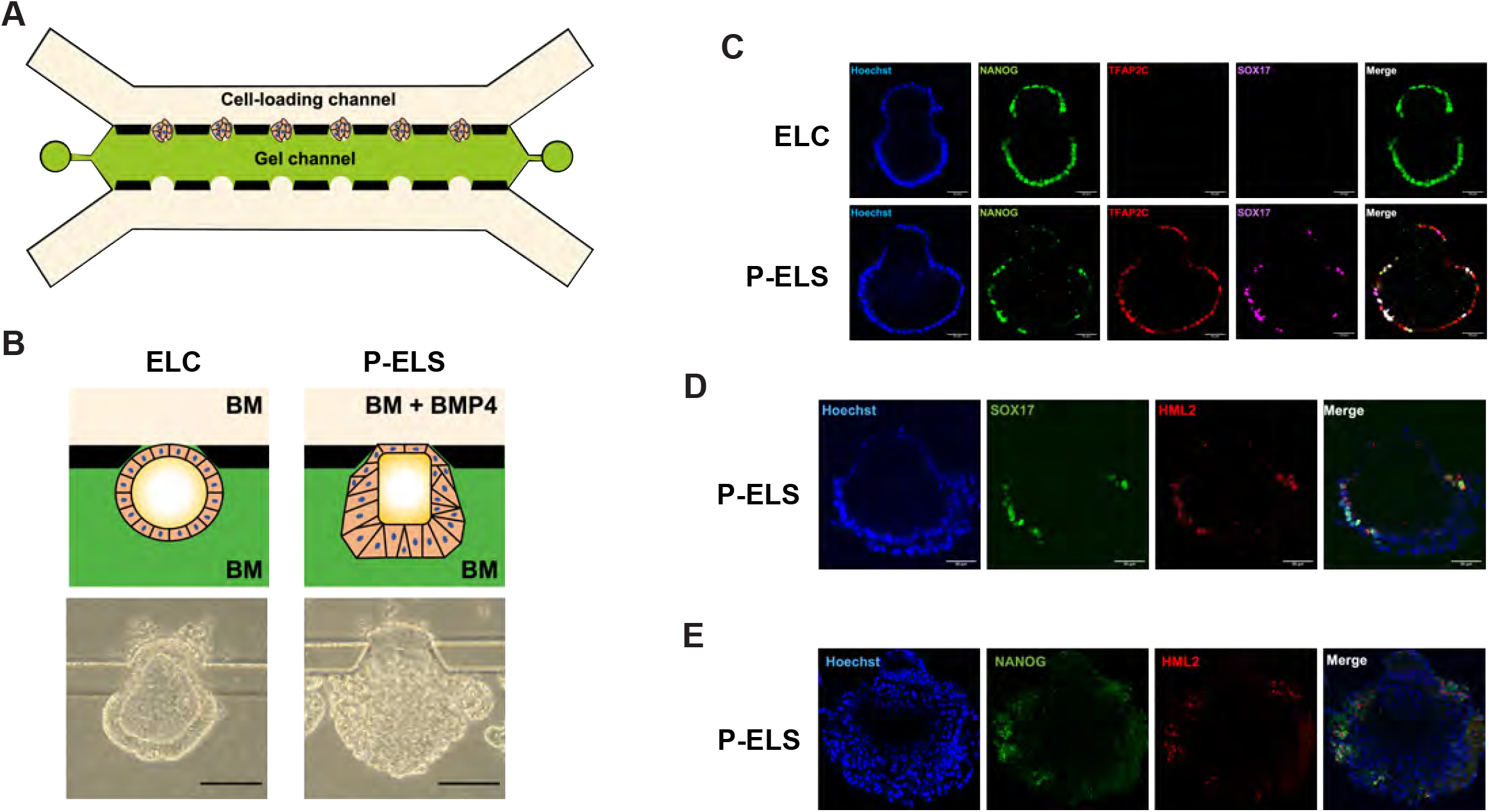
Expression of HML2 RNA in human embryoids generated with a polarized exposure to BMP4 in a microfluidics device. (A, B) Schematic representation of the microfluidics device for polarized exposure to human iPSC aggregates. (A) Formation of cell aggregates at the boundary of cell-loading and gel channels. (B) Morphological characteristics of the epiblast-like cyst (ELC), and the posteriorized embryonic-like sac (P-ELS) generated in the absence or presence of polarized exposure to BMP4. BM, basal medium. (C-E) Fluorescence confocal microscopy. (C) Immunofluorescence (IF) detection of NANOG, TFAP2C, and SOX17 proteins in ELC and P-ELS. (D) SOX17 protein (IF) and HML2 RNA (RNA-ish) detection in P-ELS. (E) RNA-ish detection of NANOG and HML2 in P-ELS.

### Production of HML-2 viral proteins and VLPs in LTC-hPGCLCs

Taking advantage of the LTC-hPGCLC cell culture technique that readily yields millions of hPGCLCs (Kobayashi et al., 2022), we attempted to detect viral proteins produced by HML2. Western blotting of total cell lysates with an antibody raised to the HML-2/HERV-K group-specific antigen (GAG) protein detected a 74-kDa band, which corresponds to the GAG precursor protein (Lee and Bieniasz, 2007), in LTC-hPGCLCs but not in hiPSCs (Fig. 5A). We also detected a protein band of the same size from cell culture supernatant of LTC-hPGCLCs but not hiPSCs.

**Fig. 5.**
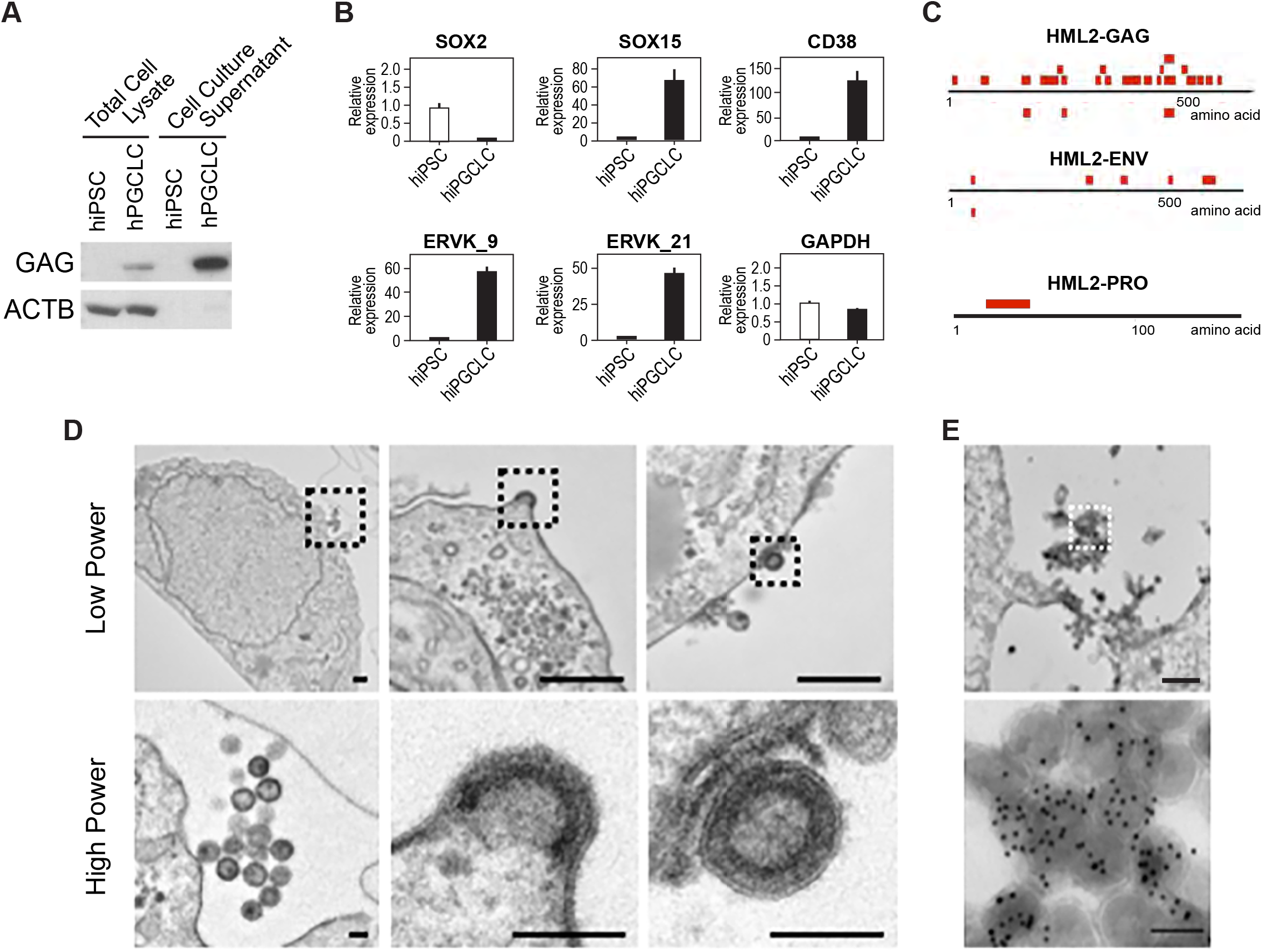
Expression of the HML2 proteins and virus-like particles (VLPs) in long-term culture human PGCLCs (LTC-hPGCLCs). (A) Western blotting detection of HML2 GAG protein in cell lysate and cell culture supernatant. ACTB, β-actin. (B) Proteomics detection of HERVK GAG proteins (HERVK_9 and HERVK_21), pluripotency marker (SOX2), and PGCLC markers (SOX15 and CD38). Bars indicate mean ± SD of triplicated measurements. (C) Locations of detected peptides in HML-2 GAG, ENV, and PRO proteins. (D, E) Transmission electron microscopy images of VLPs formed at the surface of LTC-hPGCLCs. Areas shown with dotted rectangles in the low power images are magnified in the high-power image below. Scale bars in the low and high-power images indicate 500 nm and 100 nm, respectively. (E) Immunogold staining using an anti-HERVK GAG protein demonstrates specific enrichment of the gold particles at the VLPs.

To obtain further evidence of protein expression from HML2, cell pellets of LTC-hPGCLCs and hiPSCs were subjected to quantitative proteomics analysis (Figs. 5B, 5C). SOX2 protein, a pluripotent stem cell marker, was more strongly expressed in hiPSCs than LTC-hPGCLCs whereas expression of PGCLC marker proteins SOX15 and CD38 was stronger in LTC-hPGCLCs than hiPSCs, agreeing with the mRNA expression profile reported in our preceding study (Kobayashi et al., 2022). We observed strong expression of HML2 proteins corresponding to the predicted peptides of ERVK9 and ERVK21. In LTC-hPGCLCs, multiple peptides corresponding to parts of the predicted HML2-GAG and envelope (ENV) proteins are detected. We also detected a peptide corresponding to the HML2 viral proteinase (PRO), whose translation is dependent on ribosomal frame shifts of the same RNA transcript encoding the GAG protein (Garcia-Montojo et al., 2018).

As we observed expression of HML2 proteins in LTC-hPGCLCs, we attempted to determine whether HML2 is also capable of producing VLPs. Strikingly, transmission electron microscopy readily detected VLPs on the surface of LTC-hPGCLCs (Fig. 5D) but not hiPSCs (data not shown). The VLPs were approximately 100 nm in diameter with clearly visible condensed cores and lipid bilayer surface membrane but no prominent spikes. We observed VLPs already released from the cell surface (Fig. 5D, *left* panels) as well as VLPs being formed (*center*) or still adhered (*right*) to the plasma membrane. Immunoelectron microscopy demonstrated strong labeling of the VLPs with an anti-GAG antibody conjugated with 10 nm colloidal gold particles (Fig. 5E). Taken together, our data indicate that HML2 produces not only RNA transcripts but also viral proteins and VLPs in (LTC-)hPGCLCs, suggesting that early-stage hPGCs permit retrovirus-like activities of HML2 during their normal development.

## DISCUSSION

Our current study has confirmed recent reports on expression of LTR5_Hs viral RNA in hPGCLCs (Ito et al., 2022; Xiang et al., 2022). The class switching in the dominantly active HERV species from LTR7/HERV-H to HML2 during hiPSC conversion to hPGCLC (Figs. 1, 2A, 2B, S1) agrees with our previous observation that CpG sites in LTR5_Hs were robustly demethylated along with this conversion whereas LTR7/HERV-H was demethylated only modestly (Kobayashi et al., 2022). The HML2 copies activated in hPGCLCs were characterized with LTR5_Hs flanking the protein-coding sequence whereas other HML2 copies harboring at least one non-Hs, older LTR5 were not activated (Fig. 2D), suggesting that relatively young copies of the hominoid-specific HML2 may be selectively activated in hPGCLCs. Whereas the preceding studies on LTR5_Hs activation in hPGCLCs focused on genomic effects of LTR5_Hs activation, our current study revealed that LTR5_Hs also perform retrovirus-like virological actions, including production of viral proteins or VLPs, reminiscent of the VLP production by human epiblast cells (Grow et al., 2015). It is tempting to speculate that expression of LTR5_Hs proteins may affect the innate immune system in human germline cells (Canadas et al., 2018; Chuong et al., 2016; Grandi and Tramontano, 2018; Zhao et al., 2014), which needs to be examined in future studies.

Some of the non-seminomatous human germ cell tumor cell lines derived from embryonal carcinomas/teratocarcinomas are known to produce HML2 VLPs as well as viral proteins (Bieda et al., 2001), which is reminiscent of HML2 activation in human naïve pluripotent stem cells (Grow et al., 2015). In contrast, our current study revealed that HML2 was not the HERV species predominantly activated in seminomatous human germ cell tumors, in which other HERV species such as THE1B or LTR17 were strongly activated (Figs. 1 and S1). Embryonal carcinomas express the pluripotency marker SOX2 but not the hPGC/hPGCLC marker SOX17 whereas both human seminomas and hPGCLCs are SOX2-negative and SOX17-positive (Kobayashi et al., 2022; Muller et al., 2021). It has been proposed that SOX2 and SOX17 determine the fate of germ cell tumors to either embryonic stem cell-like (embryonal carcinoma) or hPGC-like (seminoma) (Muller et al., 2021). The unique profiles of HERV activation between embryonal carcinomas and seminomas may contribute to the distinct biological characteristics of these two types of tumors derived from hPGCs.

In summary, our current study has revealed that young copies of the hominoid-specific LTR5_Hs HERVs produce not only RNA but also viral proteins and VLPs in human PGCLCs. We also provide evidence that LTR5_Hs are activated in early-stage hPGCs *in vivo* immediately after these first germline precursor cells emerge from the amnion/epiblast in the pre-gastrulation stage of human embryos. Future research on biological significance of the LTR5_Hs activation in human germ cells should study not only the genomic impact of LTR5_Hs sequences as transcriptional enhancers/activators but also potential roles of the LTR5_Hs viral proteins and VLPs in the innate and/or adaptive immunity as well as germline cell development.

## MATERIALS AND METHODS

### Human cell cultures and tissues

All human iPSCs (A4, A5, A6) used in the present research and their differentiation to hPGCLCs through microwell-supported formation of embryoid bodies were described in our previous studies (Kobayashi et al., 2022; Mitsunaga et al., 2017; Mitsunaga et al., 2021). Human seminoma tumor tissues were surgically excised from patients at the Massachusetts General Hospital (MGH) and pathologically diagnosed as pure seminomas by the MGH Genitourinary Pathology Services. Frozen tissues of the tumors were then made available for the current research through the MGH Genitourinary Tumor Bank (IRB approval number ???)

### RNA-seq

The fastq deep sequencing raw data of human iPSCs, CD38^+^ hPGCLCs, and CD38^-^ EBCs were described in our previous study (Mitsunaga et al., 2017). Total RNA extraction, library construction, and deep sequencing of human seminoma tissues were performed similarly to obtain 34-52 million, uniquely mapped paired-end reads (75 + 75).

### Quantitation of RNA expression from HERV loci

RNA-seq estimation of RNA expression from HERV loci was performed using the ERVmap Perl scripts as we previously described (Tokuyama et al., 2018). While the ERVmap tool is accessible as a web-based service (https://www.ervmap.com), we also developed a novel tool implementing the original ERVmap pipeline by Ruby scripts using updated and publicly available software tools. In the current study, this Ruby-based tool is mentioned as ERVmap2.

The FASTQ raw sequence reads were subjected to quality control analysis using the fastQC tool (Babraham Institute), and adaptor sequences, low-quality reads (Phred score < 25), and short reads (< 40 bp) were removed using the Trim Galore! tool (Babraham). The filtered FASTQ reads were either subjected to ERVmap analysis of HERV RNA expression or examined using ERVmap2 as follows. The FASTQ reads were aligned to the GRCh38/hg38 human genome reference sequence using the STAR aligner to generate BAM alignment files. Uniquely mapped reads were extracted from the BAM files using sambamba (Tarasov et al., 2015) and subjected to counting their overlaps with a BED file of HERV coordinates using bedtools (Quinlan and Hall, 2010).

Whereas the original ERVmap uses a BED file containing coordinates of 3,220 HERV proviruses, ERVmap2 uses an updated BED file containing 2,504 HERV coordinates generated using stricter criteria of proviruses. Thus, LTRs and internal HERV sequences identified in the GRCh38/hg38 human reference genome by RepeatMasker (Tarailo-Graovac and Chen, 2009) were filtered for the clade, LTR species, and internal sequence types described in Table S3. Then HERVs consisting of one 5’ LTR, one 3’ LTR, and at least one internal sequence connected via gaps not greater than 1 kb. Numbers of HERVs belonging to each clade and the whole list of the selected HERVs are provided as Tables S3 and S4, respectively. Table S4?

For evaluation of HERV RNA quantitation tools, ERVmap2 BED files of the HERV provirus list was used, and for Telescope this BED file was converted to the GTF format. FASTA sequences of the HERV proviruses were generated using bedtools. FASTQ reads simulating Illumina sequencing and the “gold standard” SAM alignment data were generated using the ART simulator (Huang et al., 2012), and the FASTQ data were subjected to analyses using HERV RNA quantitation tools ERVmap (Tokuyama et al., 2018), ERVmap2, Telescope (Bendall et al., 2019), and Salmon-TE (Jeong et al., 2018). Outcomes of the tools were compared with ERV counts generated from the gold standard SAM alignment data using bedtools. All read counts were normalized using the negative binominal trimmed mean of M-values method implemented by the Bioconductor package edgeR (Robinson et al., 2010) and inspected using by hexplot using R.

### Microfluidics-supported human embryoid formation

Microfluidic devices for formation of embryoids from hPSCs under polarized exposure to BMP4 were prepared as we previously described (Zheng et al., 2019). Human iPSCs were dissociated using Accutase (Innovative Cell Technologies, AT104) and suspended in mTeSR plus (Stemcell Technologies, 100-0276) medium containing 10μM Y27632 ROCK inhibitor (Axon Medchem, 1683) at 1.0 × 10^7^ cells/mL. The cell loading channel of the device was loaded with 1.0 × 10^5^ cells in 10 μL. To generate the posterior primitive streak-like cells, BMP4 (50 ng/mL) was added to the mTeSR Plus medium in the cell-loading channel whereas the gel channels was loaded with mTeSR Plus without BMP4.

### Immunofluorescence

Cells were fixed by 4% formaldehyde in PBS for 12 h, and permeabilized in 0.1% Triton X-100 in PBS for 1 h. After blocking in 4% donkey serum at 4°C for 3 h, cells were incubated with primary antibodies at 4 °C for 24 h and then secondary antibodies at room temperature for 6 h. Primary antibodies using in this study were goat anti-SOX17 (R&D Systems, AF1924, dilution 1:2000), rabbit anti-NANOG (Cell Signaling Technology, 4903, dilution 1:200), mouse anti-TFAP2C (Santa Cruz, sc-12762, dilution 1:200). The secondary antibodies were donkey anti-rabbit-488 (Abcam, ab150061, dilution 1:500), donkey anti-goat-568 (Abcam, ab175704, dilution 1:500), and donkey anti-mouse-647 (Abcam, ab150111, dilution 1:500). Nuclei were counter-stained using Hoechst33342 (Thermo Fisher Scientific, H21492). Fluorescence images were taken with a Zeiss LSM710 confocal microscope and processed using Image J Fiji (Schindelin et al., 2012).

### RNA *in situ* hybridization

RNA *in situ* hybridization was performed using the ViewRNA ISH Cell Assay Kit (Thermo Fisher, QVC0001) according to the manufacturer’s instructions. Embryoids developed in the microfluidic device were fixed with 4% formaldehyde for 6 h and dehydrated with a graded series of methanol (50%, 75%, and 100%) and stored. Embryoids were rehydrated using a reverse series of methanol (75%, 50% in PBS), permeabilized in 0.1% Triton X-100 in PBS for 1 h and digested with proteinase K for 10 min at room temperature. The embryoids were then hybridized with a fluorescence-labeled DNA probe targeting human HML-2 endogenous retrovirus RNA for 3 h at 40°C, followed by incubation with the preamplifier, amplifier, and label probe solutions provided in the kit for 30 min each at 40°C. Nuclei of the embryoids were counter-stained with Hoechst 33342 and visualized by fluorescence microscopy as described earlier.

### Western blotting

hiPSCs and LTC-hPGCLCs were grown in feeder-free conditions on Matrigel as we recently described (Kobayashi et al., 2022). Cell culture media were collected from subconfluent cultures ??? hours after final medium change and centrifuged at 300 x *g* for 5 min at 4 °C to remove cellular debris. Adherence cells were washed with ice-cold PBS and lysed in the RIPA buffer. Western blotting was performed as described (Kobayashi et al., 2022) using an anti-GAG (mouse monoclonal anti-GAG, AUSTRAL Biologicals, HERM-1841-5, dilution 1:10,000) and anti-β-actin (company, cat#, dilution?) primary antibodies and horseradish peroxidase-conjugated anti-mouse Ig secondary antibody (Santa Cruz, sc-516102).

### Quantitative proteomics

hPGCLC cultures derived from male hiPSC clones A4 and 9A13 were produced and expanded *in vitro* for 138 86 days, respectively, as we described (Kobayashi et al., 2022). LTC-hPGCLCs and their parental hiPSCs were washed with cold PBS and centrifuged to obtain frozen cell pellets, each of which consisted of 2.5 million cells. Quantitative proteomics detection of HERV proteins was performed as we previously described (Ebright et al., 2020). Briefly, total proteins were extracted from frozen cell pellets, and their disulfide bonds were reduced followed by alkylation of free cysteine thiols. Proteins were digested by the Lys-C and trypsin endoproteinases, labeled with the TMT reagents, and subjected to analysis via reversed phase LC-M2/MS3 on an Orbitrap Fusion mass spectrometer. Proteins from which the digested peptides were derived were estimated against a proteomics database, including the HERV-K proteins GAG, POL, ENV, REC, and PRO.

### Immunoelectron microscopy

Transmission and immunoelectron microscopy were performed as we described (Wilkie et al., 2022). Briefly, subconfluent human iPSCs (clone A4), LTC-hPGCLCs derived from them, and human NCCIT embryonal carcinoma cell lines were fixed with 4% paraformaldehyde and 0.1% glutaraldehyde in PBS and subjected to the standard transmission electron microscopy with negative staining using uranyl formate. For immunodetection of HERV-K VLPs, grids were stained with an anti-HERVK capsid mouse monoclonal antibody (AUSTRAL Biologicals, HERM-1831-5, dilution 1:30) followed by secondary staining with protein A conjugated with gold particles. The grids were examined on a JEOL 1200EX transmission electron microscope, and images were recorded with an AMT 2k CCD camera.

## Acknowledgements

We are grateful to Dr. Akiko Iwasaki and Dr. Yong Kong at Yale University School of Medicine for fruitful discussions and bioinformatics assistance. We also thank the Massachusetts General Hospital FACS core facility for technical help in collecting hPGCLCs.

## Competing interests

The authors declare no competing or financial interests.

## Author contributions

Experiments: MuK, MiK, JoK, JJK, KS, HH, JO, AN, YZ, ME; Computational analysis: MuK, EZ; Writing: MuK, MaT, TS; Supervision: JF, MeT, KK, SS, DI, WH, CLW, TS; Conception: TS

## Funding

This research was generously supported by the RICBAC Foundation and the Escher Fund for Autism gifts to TS. TS was also supported by NIH grants R01ES020454, R01ES023316, R01ES031139, and John Templeton Foundation Genetics Research Award #6228.

## Data availability

All RNA-seq data described in this study are available from Gene Expression Omnibus (accession numbers GSE102943 and GSE??????).

---

**Fig. S1:**
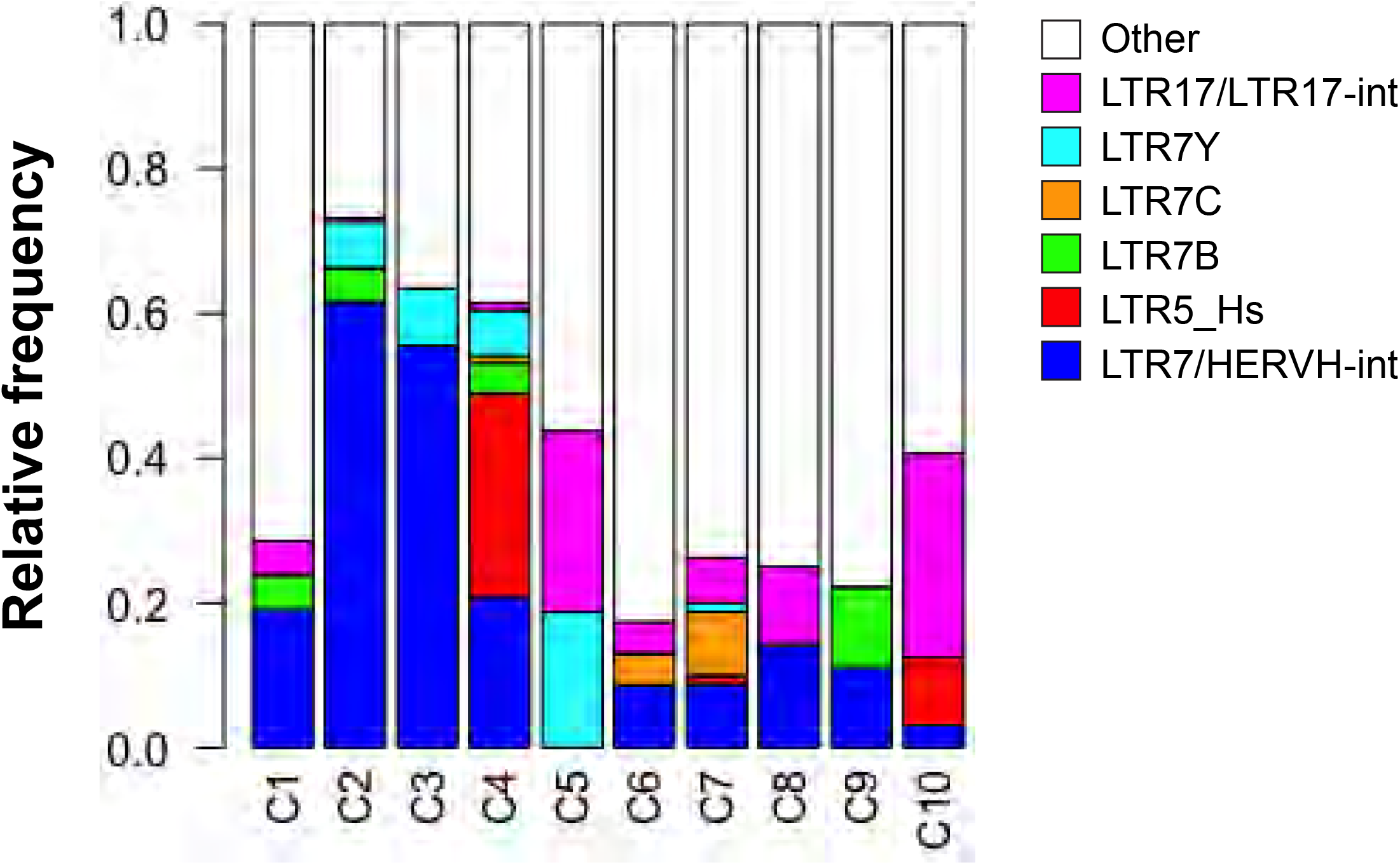
Distribution of HERV species in cell/tissue type-specific HERV clusters.

**Fig. S2:**
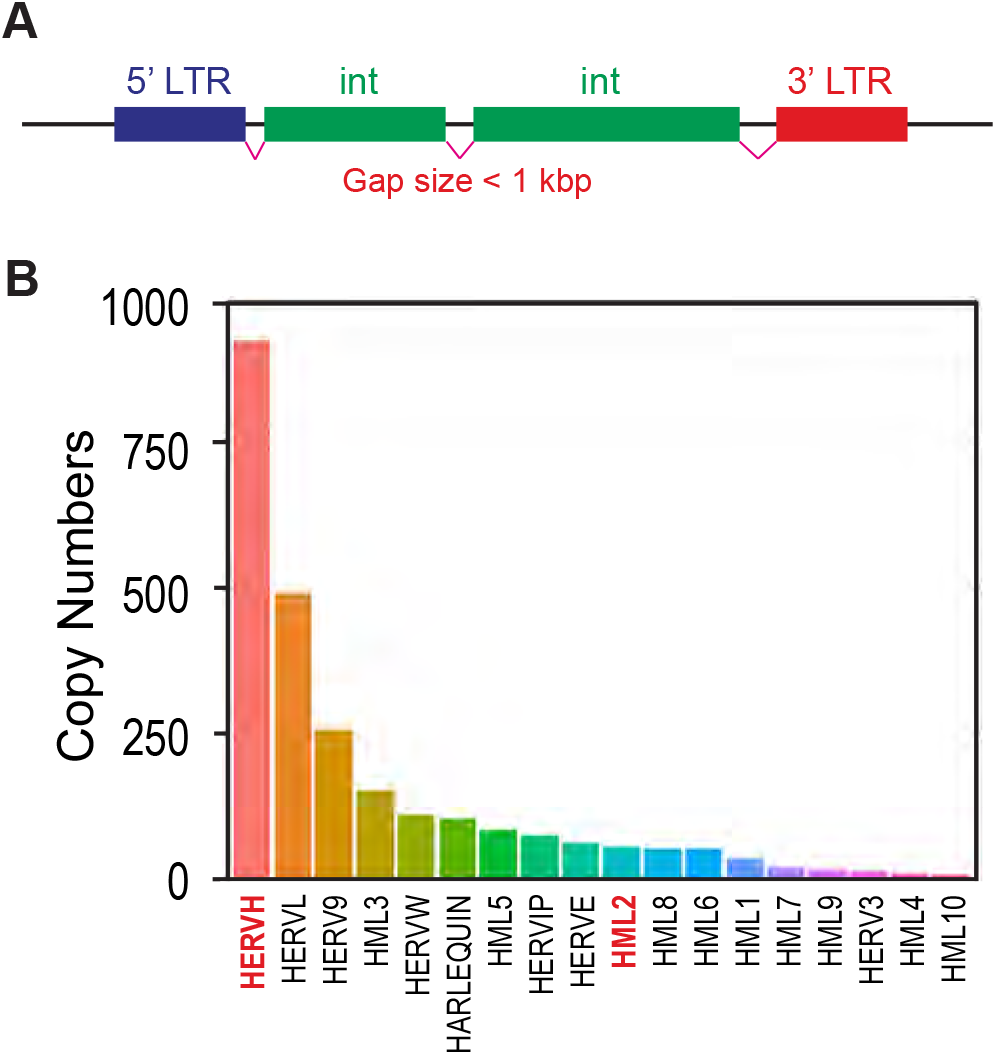
Copy numbers of well-organized HERV proviruses in the human genome. (A) Definition of a well-organized HERV provirus. (B) Copy numbers of the well-organized HERV proviruses belonging to the 18 HERV clades detected in the GRCh38/hg38 human reference genome sequence.

**Fig. S3.**
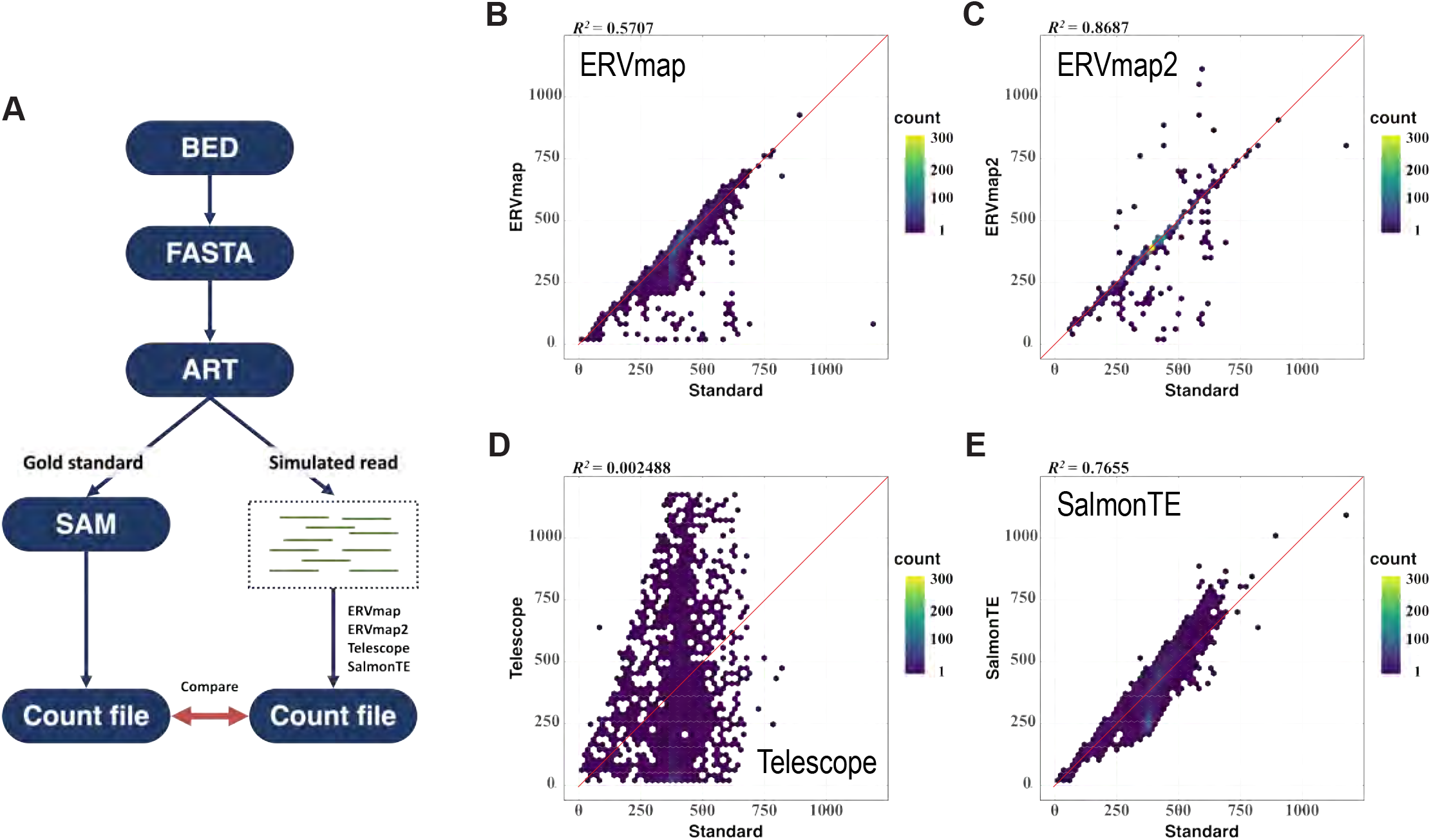
Evaluation of computational tools for determination of RNA expression from the well-organized copies of HERV proviruses. (A) Analytical scheme. FASTA format sequences of the well-organized copies of HERV proviruses were generated using coordinates of Table S2. Simulated reads resembling Illumina sequencing data (FASTQ format) and the “gold standard” read read alignment data (SAM format) were generated using the ART simulator tool. The simulated FASTQ data were subjected to HERV RNA expression analysis using (B) original ERVmap, (C), ERVmap2, (D), Telescope, and (E) SalmonTE. Panels (B-E) are hexbin plots comparing the gold standard counts (X axis) and the counts reported by each tool (Y axis). Thus, when outcomes of a tool agrees with the gold standard, datum points align along the Y=X line (red) whereas over- and underestimated HERV counts are reflected by datum points above or below the Y=X line, respectively.

**Fig. S4.**
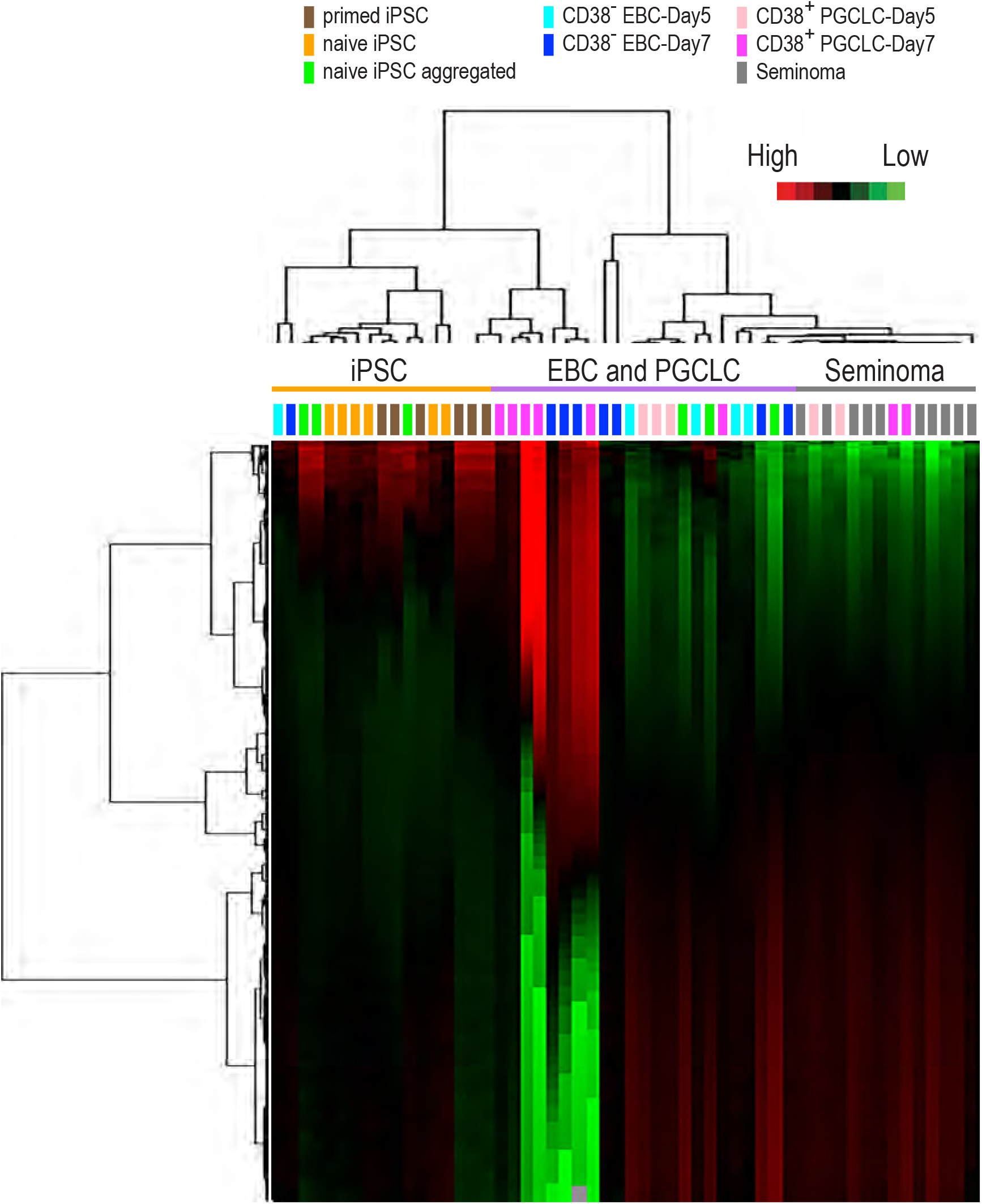
RNA-seq profiling of human iPSCs, embryoid bodies, PGCLCs, and seminoma tissues for expression of HERV RNA using Telescope. Heatmap representations of unsupervised clustering of HERV RNA expression. Color-coded cell/tissue types are shown on top of the heatmap.

**Table S3.** Copy numbers of well-organized HERVs in the 18 HERV clades in GRCh38/hg38.

| **HERV clade** | **LTR** | **Internal (int) sequence** | **Copy Numbers** |
| --- | --- | --- | --- |
| HERVE | LTR2, LTR2A, LTR2B, LTR2C | HERVE_a-int<br>HERVE-int | 61 |
| HERV3 | LTR4 | HERV3-int | 12 |
| HERVW | LTR17 | HERV17-int | 109 |
| HERV9 | LTR12, LTR12_, LTR12B, LTR12C,<br>LTR12D, LTR12E, LTR12F | HERV9-int<br>HERV9N-int<br>HERV9NC-int | 255 |
| HERVIP | LTR10B, LTR10B1, LTR10B2, LTR10F | HERVIP10B3-int<br>HERVIP10F-int<br>HERVIP10FH-int | 74 |
| **HERVH** | LTR7, LTR7A, LTR7B, LTR7Y | HERVH-int | 923 |
| HERVL | MLT2A1, MLT2A2, MLT2B3 | HERVL-int | 489 |
| HARLEQUIN | LTR2, LTR2A, LTR2B, LTR2C | Harlequin-int | 102 |
| HML1 | LTR14, LTR14A, LTR14B | HERVK14-int | 32 |
| **HML2** | LTR5, **LTR5_Hs**, LTR5A, LTR5B | HERVK-int | 55 |
| HML3 | MER9a1, MER9a2, MER9a3 | HERVK9-int | 153 |
| HML4 | LTR13, LTR13_, LTR13A | HERVK13-int | 11 |
| HML5 | LTR22, LTR22A, LTR22B, LTR22B1,<br>LTR22B2, LTR22C0, LTR22C2, LTR22E | HERVK22-int | 83 |
| HML6 | LTR3, LTR3A, LTR3B, LTR3B_ | HERVK3-int | 52 |
| HML7 | MER11D | HERVK11-int | 20 |
| HML8 | MER11A, MER11B, MER11C | HERVK11-int | 53 |
| HML9 | LTR14C | HERVK14C-int | 15 |
| HML10 | MLT14 | HERVKC4-int | 5 |

## Notes

### Competing Interest Statement

The authors have declared no competing interest.

